# Hypoosmolar dose-dependent swelling occurs in both pyramidal neurons and astrocytes in acute hippocampal slices

**DOI:** 10.1101/157180

**Authors:** Thomas R. Murphy, David Davila, Nicholas Cuvelier, Leslie R. Young, Kelli Lauderdale, Devin K. Binder, Todd A. Fiacco

## Abstract

Normal nervous system function is critically dependent on the balance of water and ions in the extracellular space. Pathological reduction in brain interstitial osmolarity results in osmotically-driven flux of water into cells, causing cellular edema which reduces the extracellular space and increases neuronal excitability and risk of seizures. Astrocytes are widely considered to be particularly susceptible to cellular edema due to selective expression of the water channel aquaporin-4 (AQP4). The apparent resistance of pyramidal neurons to osmotic swelling has been attributed to lack of functional water channels. In this study we report rapid volume changes in CA1 pyramidal cells in hypoosmolar ACSF (hACSF) that are equivalent to volume changes in astrocytes across a variety of conditions. Astrocyte and neuronal swelling was significant within 1 minute of exposure to 17 or 40% hACSF, was rapidly reversible upon return to normosmolar ACSF, and repeatable upon re-exposure to hACSF. Neuronal swelling was not an artifact of patch clamp, occurred deep in tissue, was similar at physiological vs. room temperature, and occurred in both juvenile and adult hippocampal slices. Neuronal swelling was neither inhibited by TTX, nor by antagonists of NMDA or AMPA receptors, suggesting that it was not occurring as a result of excitotoxicity. Surprisingly, genetic deletion of AQP4 did not inhibit, but rather augmented, astrocyte swelling in severe hypoosmolar conditions. Taken together, our results indicate that neurons are not osmoresistant as previously reported, and that osmotic swelling is driven by an AQP4-independent mechanism.

## 1 Introduction

Acute reduction of plasma osmolarity in humans is a medical emergency, often resulting in seizures and sometimes coma or even death (Andrew 1991; Castilla-Guerra et al. 2006). Such deleterious effects on the CNS can be attributed to the sudden change in osmotic pressure within the interstitial space of the brain, which causes cells to take on water leading to “cellular” or “cytotoxic” edema (Kimelberg 1995). Tissue swelling resulting from cellular edema has been observed in many studies examining the effect of hypoosmolar conditions on excitability (Andrew and MacVicar 1994; Chebabo et al. 1995b; Kilb et al. 2006). Given that cell swelling shrinks the extracellular space (ECS) and increases tissue resistance, hypoosmolar conditions amplify nonsynaptic excitability in neurons and increase susceptibility to seizure (Lauderdale et al. 2015; Roper et al. 1992; Rosen and Andrew 1990; Schwartzkroin et al. 1998).

Most cellular edema in the brain is thought to be driven by water influx through aquaporin-4 (AQP4) channels, which are expressed primarily by astrocytes (Nagelhus et al. 2004). Knockout of AQP4 inhibits the tissue swelling normally associated with seizures (Binder et al. 2004), stroke (Katada et al. 2014), and other models of cytotoxic edema (Papadopoulos and Verkman 2013). In normal physiology, AQP4 is important for water homeostasis in the brain, and appears to be required for the activity-dependent fluxes of water which occur alongside potassium uptake and buffering (Amiry-Moghaddam et al. 2003; Eid et al. 2005; Illarionova et al. 2010; Strohschein et al. 2011). Astrocytic swelling via AQP4-mediated water uptake is well-established as both a physiological and pathological phenomenon.

Neuronal volume changes, by contrast, are considered to be almost exclusively pathological. Neuronal somata will rapidly swell in cases of oxygen-glucose deprivation (OGD) and excitotoxic damage (Andrew et al. 2007; Choi 1992; Liang et al. 2007;Risher et al. 2009; Rungta et al. 2015), but appear very resilient to hypoosmolar swelling, an expected consequence of nonfunctional water channels (Andrew et al. 2007; Caspi et al. 2009; Gorelick et al. 2006). This has led to the conclusion that hypoosmolar conditions selectively swell glial cells (Andrew et al. 2007; Risher et al. 2009). However, this conclusion contradicts earlier reports of swelling under hypoosmolar stress in isolated neurons in culture (Aitken et al. 1998; Borgdorff et al. 2000; Somjen 1999). Furthermore, studies which have directly examined hypoosmolar swelling of neurons or astrocytes (Andrew et al. 2007; Hirrlinger et al. 2008; Risher et al. 2009) have lacked temporal resolution below approximately 5 minutes. We have recently found that neuronal excitability increases significantly within 2 minutes of exposure to hypoosmolar ACSF (hACSF), suggesting that constriction of the ECS occurs rapidly during hACSF application (Lauderdale et al. 2015). Determining the precise contribution of cellular swelling and reduction of the ECS to rapid changes in neuronal excitability requires an equally rapid measurement of cell volume.

In the present study, we used confocal imaging to quantify stratum radiatum (s.r.) astrocyte and CA1 pyramidal neuron volume in acute mouse hippocampal slices during repeated 5-7 minute hypoosmolar ACSF (hACSF) applications, at 1-minute resolution. Surprisingly, we found that CA1 neurons and s.r. astrocytes swell to the same extent and over a similar time course in both 17% and 40% hACSF. Neuronal swelling occurred across a variety of experimental conditions, suggesting that it is not an isolated phenomenon or attributable to one particular variable. Neuronal swelling did not differ between adult and juvenile animals, nor was it prevented by NMDA receptor antagonists, AMPA/Kainate receptor antagonists, TTX, or AQP4 deletion. Astrocyte swelling was also unaffected by AQP4 deletion in 17% hACSF and significantly augmented, rather than inhibited, in 40% hACSF. These data provide evidence that neurons are not osmoresistant and suggest that neurons and astrocytes may share a common, AQP4-independent swelling pathway in hypoosmolar conditions.

## 2 Methods

All animals and protocols used in the following experiments were approved by the Institutional Animal Care and Use Committee at the University of California, Riverside.

### 2.1 Slice preparation in juveniles

Hippocampal slices were prepared from juvenile (15-21 day old) C57Bl/6J mice as previously described (Xie et al. 2014). In some experiments, Thy1-eGFP (Jackson Laboratories, Bar Harbor, ME) or AQP4^-/-^ mice were utilized, both of which were on a C57Bl/6J background and exhibited no obvious phenotypic or behavioral differences from wild-type (Feng et al. 2000; Ma et al. 1997). Animals were deeply anesthetized under isoflurane and decapitated, and brains were quickly moved into ice-cold “slicing buffer” containing (in mM): 125 NaCl, 2.5 KCl, 3.8 MgCl_2_, 1.25 NaH_2_PO_4_, 26 NaHCO_3_, 25 glucose, and 1.3 ascorbic acid, bubbled continuously with carbogen (95% O_2_/5% CO_2_). Parasagittal hippocampal slices (350 μm thick) were prepared using a Leica VT1200S vibratome (Leica, Nussloch, Germany) and transferred to a recovery chamber containing standard artificial cerebrospinal fluid (ACSF), which was composed of (in mM): 125 mM NaCl, 2.5 KCl, 2.5 CaCl_2_, 1.3 mM MgCl_2_, 1.25 NaH_2_PO_4_, 26 NaHCO_3_, and 15 glucose, bubbled continuously with carbogen (~299-303 mOsm). Slices were incubated in ACSF at 36° C for 45 minutes, then allowed to cool to room temperature for a minimum of 15 minutes before being transferred to a recording chamber for experiments.

In some cases, sulforhodamine-101 (SR-101; Sigma-Aldrich, St. Louis, MO) was used during incubation to selectively stain astrocytes (Schnell et al. 2015). Incubation was adjusted as follows (Xie et al. 2014): After preparation, slices were transferred to a recovery chamber containing 1 μM SR-101, dissolved in a modified ACSF composed of (in mM): 125 NaCl, 2.5 KCl, 0.5 CaCl_2_, 6 MgCl_2_, 1.25 NaH_2_PO4, 26 NaHCO3, 15 glucose and 1.3 ascorbic acid, bubbled continuously with carbogen, and incubated at 36° C for 25-35 minutes. Slices were subsequently transferred to the same modified ACSF without SR-101 for 10 more minutes at 36° C. Finally, slices were allowed to cool to room temperature for 15 minutes before being transferred to ACSF. Slices were equilibrated in ACSF for at least 15 minutes prior to use. For experiments performed at physiological temperature, slices were kept at 36° C.

### 2.2 Slice preparation in adults

Adult (10-15 week old) hippocampal slices were prepared from C57Bl/6J or Thy1-eGFP mice as previously described (Lauderdale et al. 2015). Adult slicing buffer (ASB) contained (in mM): 87 NaCl, 75 sucrose, 2.5 KCl, 0.5 CaCl_2_, 7 MgCl_2_, 1.25 NaH_2_PO_4_, 25 NaHCO_3_, 10 glucose, 1.3 ascorbic acid, 2 pyruvate, 3.5 MOPS, and 100 μM kynurenic acid, bubbled continuously with carbogen, and was partially frozen to form an icy “slush”. Slices were prepared as described above, and transferred to a recovery chamber containing carbogen-bubbled ASB at 36° C for 45 minutes. Following incubation, slices were allowed to cool to room temperature for 15 minutes before being transferred to standard ACSF, where they equilibrated for a minimum of 20 minutes before being used for experiments.

The SR-101 protocol was adapted for use in adults by adding SR-101 to the ASB which was similar in composition to the modified ACSF normally used for SR-101 loading. For unknown reasons, adult astrocytes proved far more resistant to SR-101 loading than did juvenile astrocytes, and required additional modifications to our labeling protocol. We found that a 45-minute incubation in 2-4 μM SR-101 labeled adult astrocytes to a similar degree and depth as juvenile astrocytes, with limited background staining. Slices were cooled to room temperature for 15 minutes in SR-101-free ASB after incubation, and then transferred to standard ACSF as above.

### 2.3 Solutions and drugs

“Normosmolar ACSF” (nACSF, ~298 mOsm), consisting of standard ACSF without MgCl_2_, was used for baseline recordings and wash periods. Seventeen or 40% “hypoosmolar ACSF” (hACSF) was prepared by using deionized water to dilute nACSF by 10% (final osmolarity: ~270 mOsm), 17% (final osmolarity: ~250 mOsm) or 40% (final osmolarity: ~180 mOsm). These percent reductions in osmolarity have been generally described as “mild” to “modest” (< 20% hypoosmolarity) (Anderova et al. 2014; Azouz et al. 1997; Thrane et al. 2011) or “moderate” to “severe” (> 30% hypoosmolarity) (Anderova et al. 2014; Kimelberg 2004; Kimelberg et al. 2006), and were chosen based on those often used in previous studies which have ranged anywhere from 5% to 65% (Andrew et al. 2007; Chebabo et al. 1995b; Darby et al. 2003; Fiacco et al. 2007; Haskew-Layton et al. 2008; Hirrlinger et al. 2008; Kilb et al. 2006; Kimelberg 2004; Lauderdale et al. 2015; Thrane et al. 2011; Wurm et al. 2010). Unless otherwise indicated, “experimental” ACSF solutions (nACSF and hACSF) contained 1 μM TTX (Cayman Chemical, Ann Arbor, MI) to block voltage-gated Na+ channels, and 10 μM NBQX (Alomone Labs, Jerusalem, Israel) to block AMPA/Kainate receptors (Lauderdale et al. 2015). In some experiments, 50 μM AP5 (an NMDA-receptor antagonist; Abcam, Cambridge, MA) or 6 mM Mg^2+^ (to maximize block of NMDA receptors in the slice; Dissing-Olesen et al. 2014) were also added to the experimental solutions.

A number of experiments were performed using more “physiological” antagonist-free solutions, which were based on standard (1.3 mM Mg^2+^) ACSF and contained no TTX or NBQX. In these experiments, standard ACSF was used in place of nACSF for baseline and wash periods, and diluted by 17% or 40% to generate the respective hACSF doses.

### 2.4 Patch-clamp of neurons and astrocytes

Hippocampal slices were transferred to a recording chamber and continuously superfused with room-temperature, oxygenated ACSF. For experiments at physiological temperature, recording chamber and perfusion solution temperatures were maintained at 36°C using a Warner Instruments TC-344B Dual Channel Heater Controller (Warner Instruments, Hamden, CT). Slices were visualized using an Olympus BX61 WI upright microscope equipped with UMPLFLN 10x (N.A. 0.3) and LUMPlanFl 60x (N.A. 0.9) water-immersion objectives and DIC optics (Olympus America, Center Valley, PA). CA1 pyramidal neurons and passive stratum radiatum (s.r.) astrocytes were patch clamped in order to load them with fluorescent indicator dyes for volume measurements in our initial experiments. Whole-cell patch clamp was performed using a Multiclamp 700B amplifier and Digidata 1550 digitizer, controlled through pClamp v.10.4 and Multiclamp commander software (Molecular Devices, Sunnyvale, CA). Patch pipettes were pulled from borosilicate glass using a Narishige PC-10 vertical micropipette puller (Narishige, Tokyo, Japan). Neuronal patch pipettes had a resistance of 3.8-5.3 MΩ when filled with an internal solution containing (in mM): 140 K-gluconate, 4 MgCl_2_, 0.4 EGTA, 4 Mg-ATP, 0.2 Na-GTP, 10 HEPES, and 10 phosphocreatine, pH 7.3 with KOH. For dye loading, neuronal patch pipettes also contained the fluorophores Alexa 488 hydrazide (100-200 μm) or Alexa 594 hydrazide (200 μM; Thermo Fisher Scientific, Waltham, MA). Astrocyte patch pipettes had resistances of 4.2-8.9 MΩ when filled with an internal solution containing (in mM): 130 K-gluconate, 4 MgCl_2_, 10 HEPES, 10 glucose, 1.185 Mg-ATP, 10.55 phosphocreatine, and 0.1315 mg/ml creatine phosphokinase, pH 7.3 by KOH. Astrocyte pipettes also contained Alexa Fluor 488 dextran or Oregon Green 488 dextran (Thermo Fisher Scientific), both 10 kDa in size to prevent dye spread into neighboring astrocytes via gap junctions.

Neurons and astrocytes were identified first by location and morphology under DIC optics, and confirmed by their electrophysiological properties. Pyramidal neuron resting V_m_ was-60.1 ± 0.7 mV (n = 30) and exhibited characteristic voltage-gated Na^+^ and K^+^ currents in response to a voltage step protocol. Astrocytes were identified as “passive” if they exhibited a characteristically low input resistance, low resting membrane potential (-78.0 ± 1.2 mV; n = 15) and lack of voltage-gated conductances. Astrocytes and neurons were voltage-clamped to-90 or-70 mV, respectively, for no more than 5 minutes to allow for dye diffusion into the cytoplasm (with rare exceptions for dextran-loading of astrocytes, which sometimes required up to 8 minutes). In the interest of limiting the amount of cytoplasm dialyzed by the internal solution, this time was kept to a minimum with occasional, quick confocal scans to check cell brightness. Once dye loading was deemed sufficient for imaging, the pipette was gently withdrawn. A smooth, stable “off-cell” and formation of a > 1GΩ seal during pipette removal was considered an indicator that the cell was not damaged during withdrawal of the patch pipette. All patch clamped cells were allowed to recover for at least 10 minutes before further use. In later experiments, patch clamp was mostly supplanted by bulk loading astrocytes with SR-101 dye (see above), and by using neurons from the Tg(Thy1-EGFP)MJrs/J (Thy1-GFP-M, stock #7788) or B6;CBA-Tg(Thy1-EGFP)SJrs/NdivJ (Thy1-GFP-S, stock #11070) mouse lines, which express eGFP under the neuronal Thy1 promoter in some pyramidal neuron populations (Feng et al. 2000). In these instances, cells were chosen based on their depth in the tissue, lack of obvious morphological abnormalities, and their visibility under our standard imaging settings (see below).

### 2.5 Confocal imaging settings and experimental design

Alexa Fluor 488 dextran, Oregon green 488 dextran, Alexa Fluor 488, and eGFP were excited using a 488 nm argon laser (Melles Griot, Carlsbad, CA) and detected with a 503-548 nm bandpass filter, controlled by Olympus Fluoview 1000 software. Laser power was generally held at ≤ 2.0%, well below the level needed to induce photobleaching (> 50%). Pixel dwell time was 8 μs/pixel for astrocytes (which often required extra exposure time due to limited dextran loading) and 4 μs/pixel for neurons.

Alexa Fluor 594 and SR-101 were excited using a 559 nm semiconductor laser and detected using a 624-724 nm bandpass filter. Pixel dwell times were kept the same as above for consistency. Laser power ≤ 1.5% was sufficient to detect SR-101 labeled astrocytes and Alexa Fluor 594 labeled neurons. To strike an appropriate balance between image resolution and brightness, confocal aperture size was set to 300 μm and PMT voltage ~830 V across all experiments.

In one experiment, Thy1-eGFP neurons were examined much deeper within the slice (>60 μm below slice surface) and were in many cases impossible to image using our standard settings. Instead, laser power was increased to 10% and pixel dwell time to 8 μs/pixel, increasing acquisition time per image stack (~15-30 seconds) but significantly boosting cell visibility. We observed no deleterious effects on cell health resulting from the increase in laser power or exposure time.

All experiments started in standard ACSF. Where applicable, standard ACSF was replaced by nACSF following patch pipette removal or identification of the cell to be imaged, and allowed to wash in for 10 minutes prior to imaging. Imaging consisted of confocal z-stacks taken through the cell soma, beginning with a single “baseline” stack. As observed by other groups (Hirrlinger et al. 2008; Risher et al. 2009), we found that single images were insufficient for gathering the full extent of the cell body in the x-y plane (which is necessary for soma area measurements, our proxy for cell volume), and were particularly vulnerable to swelling-induced z-shifts across time points. We instead opted for rapid z-stacks through the soma, at 1.0 μm intervals and a zoom level of 3.5x (0.118 μm/pixel). X, Y and Z-shifts were compensated using quick confocal scans to check cell position, and adjusting X-Y position and Z-scan upper/lower limits accordingly before acquiring each stack. To increase scan speed, the “clip scan” function was used to crop the scan window close around the soma, reducing acquisition time to ~15 seconds per stack. These settings allowed us to obtain full z-stacks through the cell soma at 1-minute intervals. For simplicity, a z-stack encompassing the cell soma will hereafter be referred to as a “stack”.

After a baseline stack was obtained, hACSF was applied for 5 minutes and a single stack was acquired at the end of each minute. HACSF was then “washed out” by re-application of normosmolar ACSF (either nACSF or standard ACSF) for 5 minutes, at the end of which an additional stack was acquired. This sequence was repeated an additional 2 times to determine repeatability of effects on cell volume. Each subsequent hACSF application and wash period was lengthened by 1 minute (after which an additional stack was acquired), due to our observations (unpublished) that neuronal slow inward currents (SICs) evoked in these conditions were more “spread out” during subsequent hACSF applications.

### 2.6 Volume analysis protocol

Analysis of neuron and astrocyte volume was performed using the FIJI distribution of ImageJ as previously described (Lauderdale et al. 2015; Schindelin et al. 2012). Our basic analysis protocol is depicted in Figure 1. First, z-shifts across time points were corrected by choosing a common landmark and matching its slice number across all stacks. Stacks were then concatenated into an x-y-z-t hyperstack (Figure 1B), and filtered to remove noise (median filter, 2 pixel diameter). A max-intensity z-projection (MIP) was made through this hyperstack, producing a 2D time series (Figure 1C), in which each frame was a MIP showing the full x-y extent of the soma at a given time point. X-Y shifts over time were corrected using the “Linear Stack Alignment with SIFT” plugin (followed by cropping, to remove the blank areas left by image alignment), and background subtraction was performed using FIJI’s background subtraction tool (radius 50 pixels, sliding paraboloid method). The resultant time series (Figure 1D) was then binarized using the “mean” thresholding algorithm (Figure 1E) and an elliptical ROI was drawn to narrowly encompass the soma across all time points in the series (dashed red circle, Figure 1E). Area above threshold within this ROI (a measure of soma area) was used as a proxy for soma volume. Volume changes at a given time point are reported as percent change from baseline soma volume. Some experiments also include the “average percent change” from baseline over 3 hACSF applications. In these cases, percent change in the 2^nd^ and 3^rd^ hACSF periods is calculated based on the preceding nACSF wash instead of the original baseline, providing a more accurate measure of acute, relative volume changes during hACSF application. Representative images, unless otherwise noted, depict the thresholded images used for analysis.

**Figure 1:**
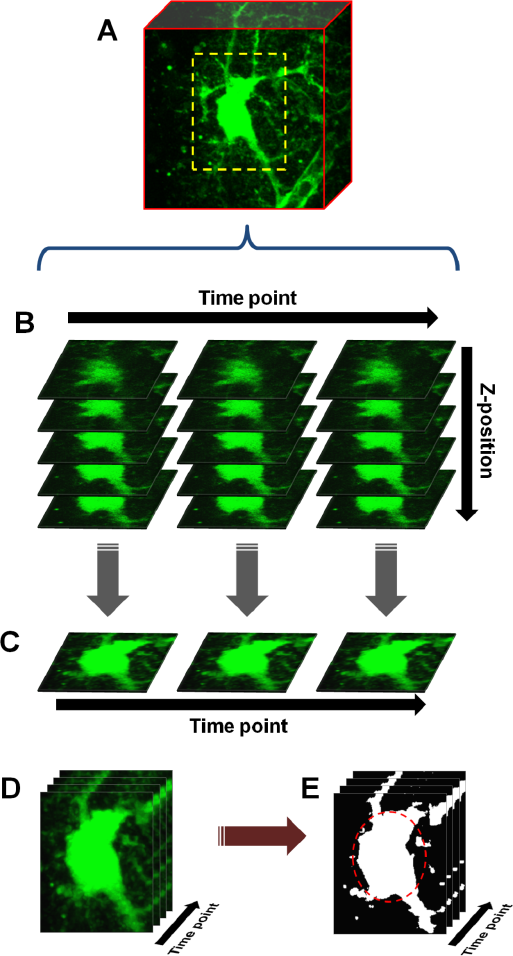
Basic protocol for cell volume analysis. (**A**) An astrocyte or neuron, fluorescently labeled by patch clamp or other means, is chosen and a scanning region (dashed yellow box) selected which closely encompasses the soma. Starting with baseline (measurement taken in normosmolar ACSF prior to hACSF application), z-stacks are taken through the soma at one minute intervals. During analysis, these stacks (x-y-z) are concatenated into a time-series (x-y-z-t) hyperstack (**B**). The hyperstack is filtered and collapsed in the z-axis to produce a time series (x-y-t) composed of max-intensity projections (**C**). After alignment, cropping and background subtraction are performed on this series, the resulting “processed” time series (**D**) is finally thresholded to produce a binary image series (**E**). An elliptical ROI (dashed red oval) is drawn around the soma, and any pixels above threshold within this ROI are quantified.

In an effort to better analyze neurons from “deep” tissue (which were not amenable to our standard analysis protocol due to poor resolution of cell borders), we employed a simplified version of the microspectrofluorimetric method of measuring volume (Crowe et al. 1995). This method does not rely on cell borders but rather on fluorescence intensity, allowing for estimation of cell volume changes using a region of interest (ROI) placed in the cell center. In brief, this method assumes a fixed number of fluorescent molecules within a cell, and therefore that any changes in cell water content (i.e. cell swelling or shrinking) will alter their concentration. As dye becomes less concentrated, fluorescence *(**F**)* should decrease linearly, and vice versa. Cell volume changes are thus inversely related to changes in cell fluorescence 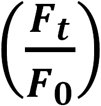, where ***F_t_*** = cell fluorescence at the time point of interest and ***F_0_*** = baseline cell fluorescence), and can be roughly calculated as the inverse of the fluorescence ratio 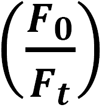. Fluorescence ratios were determined by drawing an ROI 5 μm = 5 μm wide over the center of a given cell in a processed MIP time series stack (just prior to thresholding) and measuring average intensity in this region at each time point.

### 2.7 Statistical Analysis

Statistical analysis was performed with SPSS Statistics 22 or 23 software (IBM Corporation, Armonk, NY). Mixed-design ANOVA was used for all tests to determine changes from baseline over time (within-subject factor) and between treatments, genotypes or cell types (between-subject factor). Outliers were in rare cases excluded from analysis, and only one outlier was removed from any particular group. A cell identified as an outlier was considered for removal only if: (a) its removal improved the reliability of ANOVA results (making group N more equal, reducing normality or homogeneity of variance violations, etc); and (b) measurement errors were clearly responsible for the aberrant data. Significant within-subject effects were investigated further by pairwise comparisons of each time point group to baseline. Multiple comparisons between time points were adjusted performed using the Holm-Bonferroni method, a stepwise procedure with the same assumptions as the Bonferroni correction but substantially more power for larger numbers of comparisons. Significant between-subject effects were further investigated by Student’s t-test or (if 3 or more groups) using pairwise comparisons with Bonferroni correction. Significant interactions were investigated in two steps. First, between-group simple effects were tested at each time point using Student’s t-test (for 2 groups) or one-way ANOVA followed by Tukey’s HSD post-hoc tests (for 3 or more groups). In the case of normality or homoscedasticity violations, appropriate alternatives were chosen (e.g. Welch t-test or Mann-Whitney U). Second, within-subjects simple effects were determined by splitting the file by group and running each as an individual one-way repeated measures ANOVA, with Holm-Bonferroni post-hoc tests as above. This “split-file” method has the effect of splitting the error terms by group and was deemed to be more accurate than obtaining simple main effects within the original mixed ANOVA, as the latter uses a pooled error in its calculations. N = 7-10 cells per group (after outlier removal) for all experiments, unless indicated otherwise. Significant differences are reported at the p < 0.05 (*), p < 0.01 (**), and p < 0.001 (***) levels.

## 3 Results

In a previous study, we found heightened neuronal excitability within one minute of exposing hippocampal slices to 17% or 40% hypoosmolar ACSF (Lauderdale et al. 2015). The primary focus of the current study was to examine the volume changes exhibited by neural cells over this time period. While it has been reported that hypoosmolar volume changes are mainly limited to astrocytes (Andrew et al. 2007; Caspi et al. 2009; Risher et al. 2009), conflicting reports suggest this may not always be the case (Aitken et al. 1998; Borgdorff et al. 2000; Boss et al. 2013). Therefore, we chose to examine volume responses of stratum radiatum (s.r.) astrocytes as well as CA1 pyramidal neurons, using hypoosmolar solutions (denoted hACSF for simplicity) made by 17% or 40% dilution of standard ACSF with distilled water, and including TTX (1 μM), NBQX (10 μM) and 0 mM Mg^2+^ by default. These conditions were chosen to match our previous work demonstrating significant stimulation of NMDA receptor currents by hACSF in CA1 pyramidal neurons (Lauderdale et al. 2015). The use of TTX to block neuronal firing also reduced the likelihood of spreading depression, a wave of depolarization and subsequent synaptic silencing which has sometimes been observed in hypoosmolar conditions (Chebabo et al. 1995a). Normosmolar ACSF (nACSF), used during the baseline and wash periods, was not diluted but otherwise identical in composition to the above.

### 3.1 Astrocytes and neurons swell to approximately equal volumes in hypoosmolar ACSF

We first examined the volume responses of CA1 pyramidal neurons and s.r. astrocytes to repeated applications of 17% and 40% hACSF (Figure 2). Neurons (Figure 2A-D) and astrocytes (Figure 2E, F) were initially labeled by patch-clamp dialysis of the fluorescent indicators Alexa Fluor 568 hydrazide or fluorescein dextran, respectively, and imaged using confocal microscopy (see Methods). As in our previous study, image stacks through cell somata were collected at 1-minute intervals starting with a baseline in nACSF (Figure 2A1, E1), and proceeding through 5 minutes of hACSF (Figure 2A2, E2), with a final image stack acquired after a 5 minute “wash” in nACSF (Figure 2A3, E3). Stacks were post-processed and thresholded (Figure 2B to quantify percent change from baseline at each time point (Figure 1; also see methods). In contrast to some previous studies in which neuronal volume was insensitive to hypoosmolar shifts (Andrew et al. 2007; Caspi et al. 2009), we found that neurons rapidly swelled in response to both 17% and 40% hACSF, and this swelling was similar in magnitude and time course to volume changes in astrocytes (Figure 2A-D). Neuronal soma area significantly increased above baseline within 1 minute in 17% (2.04 ± 0.33%, p < 0.001) or 40% (2.09 ± 0.72%, p = 0.044) hACSF, and continued to gradually increase to a maximum of 4.72 ± 0.41% in 17% hACSF and 10.51 ± 0.93% in 40% hACSF (Figure 2D). Overlays of baseline and 5 minute 40% hACSF time points (Figure 2C2) facilitate visualization of these volume increases as magenta expansions along the outer border of the cell. Neurons largely recovered to baseline volume after a 5 minute wash period in nACSF (1.27 ± 0.39% above baseline in 17%, 2.70 ± 0.92% in 40% hACSF, p < 0.05 each), as shown in Figure 2C3 overlays. These data bore a remarkable similarity to our observations of s.r. astrocytes in the same conditions (Figure 2E, F). As reported in Lauderdale et al. (2015), astrocytes also rapidly swelled in both concentrations of hACSF within the first minute (p < 0.01) and in a dose-dependent manner. Unlike neurons, astrocytes fully recovered to baseline volume upon return to nACSF for 5 minutes.

**Figure 2:**
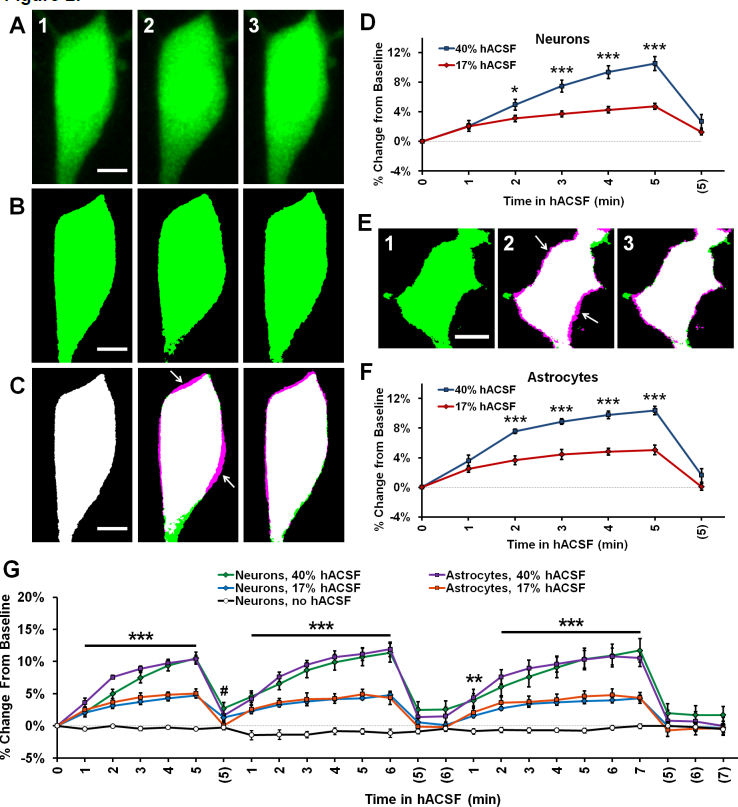
Neurons and astrocytes both swell in hypoosmolar conditions. (**A**) Representative max-intensity projections of a CA1 pyramidal neuron loaded with Alexa Fluor 488 dye via patch clamp, depicting soma volume at baseline (A1), followed by 5 minutes in 40% hACSF (A2), and a subsequent 5 minute wash period in nACSF (A3). Swelling of the soma during hACSF application is readily apparent in A2. (**B**) Time points in (A1-A3) are processed and binarized (see figure 1) to allow measurement of soma area. Images are pseudocolored green for easier comparison with (A). (**C**) Each thresholded image in (B) is pseudocolored magenta and overlaid with green baseline image from (B) to reveal regions of soma expansion during 40% hACSF, which recover to near basal levels upon return to nACSF (magenta highlights, indicated by white arrows). Scale bars = 5 μm. (**D**) Quantification of neuronal soma volume as a percent change from baseline area in 17% and 40% hACSF conditions over a single 5-minute application, followed by a 5 minute wash period denoted with (5). Time points 0, 5 and (5) correspond to images in A1, A2 and A3 respectively. (**E**) Representative thresholded images of an astrocyte loaded with Alexa Fluor 488 dextran (10,000 MW) at baseline (C1), after 5 minutes in 40% hACSF (C2), and after a 5 minute wash in nACSF (C3). As in (A), images in (C2) and (C3) have been overlaid with the baseline image to illustrate changes in cell volume (white arrows). Scale bar = 5 μm. (**F**) Quantification of astrocyte soma volume as a percent change from baseline area in 17% and 40% hACSF conditions. *p < 0.05 and ***p < 0.001 for 17% vs. 40% at each time point in B and D. All time points for both 17% and 40% hACSF were significantly elevated over baseline. (**G**) Expanded time course showing neuron and astrocyte volume changes over multiple re-applications of 40% or 17% hACSF. Neuron volume in control conditions ("no hACSF”, black line) is included for comparison. ***p < 0.001, **p<0.01 (both cell types) and #p < 0.05 (astrocytes only), percent change versus 0% (baseline). N=7-9 cells per hACSF group and 6 cells for control.

Swelling of both neurons and astrocytes was repeatable upon additional applications of 40% or 17% hACSF (Figure 2G). No neuronal swelling was detected in the absence of hACSF (Figure 2G; n = 6). No significant interaction was found between cell type and time point (40% hACSF, F(2.66,34.64) = 1.12, p = 0.351; 17% hACSF, *F*(4.11,65.73) = 1.12, p = 0.355), suggesting largely identical swelling characteristics between the two cell types.

### 3.2 Neuronal health is unlikely to be a contributing factor to hACSF-induced swelling

We began to examine the key differences in experimental protocol between our study and previous work, to determine if an unforeseen factor was responsible for (or contributing to) the neuronal swelling observed in our conditions.

Although we had no reason to suspect damage to the cell using whole-cell patch clamp approaches (see Methods), we considered the possibility that the simple act of patch clamping neurons may make them more susceptible to hACSF-induced swelling. We tested this by using hippocampal slices from transgenic Thy1-eGFP mice (Feng et al. 2000). These mice express enhanced green fluorescent protein (eGFP) in a sparse, random population of pyramidal neurons (Figure 3A), making them ideal for isolating a single neuron in the densely packed pyramidal layer of the hippocampal CA1 region (Andrew et al. 2007). Neurons expressing eGFP were bright and easily identified at our standard imaging depth of 30-40 μm (Figure 3B1, left) and once again exhibited distinct volume changes upon exposure to 40% hACSF (Figure 3B2, right). Compared to patch-loaded neurons, Thy1-eGFP neurons at this depth showed no significant difference in swelling responses to 40% hACSF (F(1,16) = 0.98, p = 0.336; Figure 3C) or 17% hACSF (F(1,16) = 1.36, p = 0.26; data not shown).

With a moderate increase in laser power (10% of maximum), we were able to extend our imaging range to ~65-90 μm below the slice surface (Figure 4A1), considerably deeper than what was feasible by patch clamp. Deeper neurons initially appeared to be more resistant to volume change in 40% hACSF (Figure 4B), with an average relative increase of only 4.22 ± 0.39% after 5 minutes in hACSF (Figure 4B, inset). This apparent reduction in osmosensitivity, however, was most likely a detection issue. Resolution of the boundary of these cells was poor, which tended to be exacerbated by the thresholding step of our analysis. For shallower neurons, both the original image (Figure 4C1) and its thresholded version (indicated by magenta outlines on Figure 4C1) tended to reflect similar cell borders. By contrast, much of the visible cell border in deeper neurons fell below threshold, and the final thresholded image did not fully represent the cell boundary (Figure 4C2). As the differences in cell volume are reflected in expansion of the cell borders, such thresholding problems would likely result in underreporting of increases in cell volume and may explain the apparent reduction in volume change by deeper neurons.

In further support of this view, we observed that both shallow (Figure 4D) and deeper neurons (Figure 4E) consistently exhibited a visible, generally similar dimming of eGFP fluorescence during 40% hACSF application (compare average intensity change between (D) and (E)). This was a strong indicator that neurons at both depths were swelling to similar degrees, as demonstrated by the microspectrofluorimetric method of volume analysis used by Crowe and colleagues (Crowe et al. 1995). Broadly speaking, if intracellular dye content is constant (as can be assumed for eGFP molecules in a typical cell), then changes in intracellular water will alter dye concentration, which can be detected via changes in fluorescence intensity. Changes in fluorescence intensity (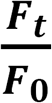, where ***F_0_*** is baseline fluorescence and ***F_t_*** the fluorescence at a given time point) are inversely proportional to changes in cell volume (Crowe et al. 1995). We reasoned that this method should be much less sensitive to detection problems than our thresholding technique (magenta outlines, Figure 4D,E), since it would not be affected by the ability to resolve the cell boundary. To quantify the eGFP intensity shift, we used the pre-thresholding MIP time series for each cell (see Methods) to examine the intensity values of a 5x5 μm region near the center of the cell (Yellow boxes, Figure 4D,E). Taking the reciprocal of the fluorescence ratio 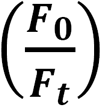, we calculated a volume increase in deep neurons of over 20% by 5 minutes in 40% hACSF (123 ± 5.36% of baseline, n = 8 cells; Figure 4F). Performing a similar measurement on neurons at our standard imaging depth (with the exception of two neurons in which eGFP was saturated and no *F_0_/F_t_* ratio could be obtained), we calculated a volume change nearly identical to that calculated in deep neurons (121 ± 3.14% of baseline). It should be noted that although the ***F_0_/F_t_*** method of volume analysis has been previously used in slices (Risher et al. 2009), its accuracy in culture is much better established (Benfenati et al. 2011; Tuz et al. 2001), and the percentages calculated above should be considered only rough approximations of the true volume change. These data do, however, provide supportive evidence that *relative* volume changes in neurons exposed to 40% hACSF are independent of depth within the slice.

We also compared patch clamp-loaded astrocytes against those loaded with sulforhodamine-101 (SR-101), a dye selectively taken up by astrocytes in the hippocampus (Schnell et al. 2012). SR-101-labeled astrocytes did not differ from patch-clamped astrocytes in their volume increase during 40% hACSF application (SR-101, n = 10; Patched, n = 7; p = 0.401; data not shown), although with repeated applications of hACSF they did appear to swell more than Thy1-eGFP neurons in the same conditions (in contrast with patch clamped astrocytes and neurons, which had shown no difference at either dose of hACSF). This effect was attributed largely to a slightly slower return to baseline in SR-101-labeled astrocytes, which caused a gradual “build up” of astrocyte volume despite no change in average rate or degree of swelling. These tests overall provided clear evidence that neuronal swelling was not an artifact of neuronal damage due to whole-cell dialysis with fluorescent indicator, nor was it likely a result of the depth at which neurons resided in the tissue.

**Figure 3:**
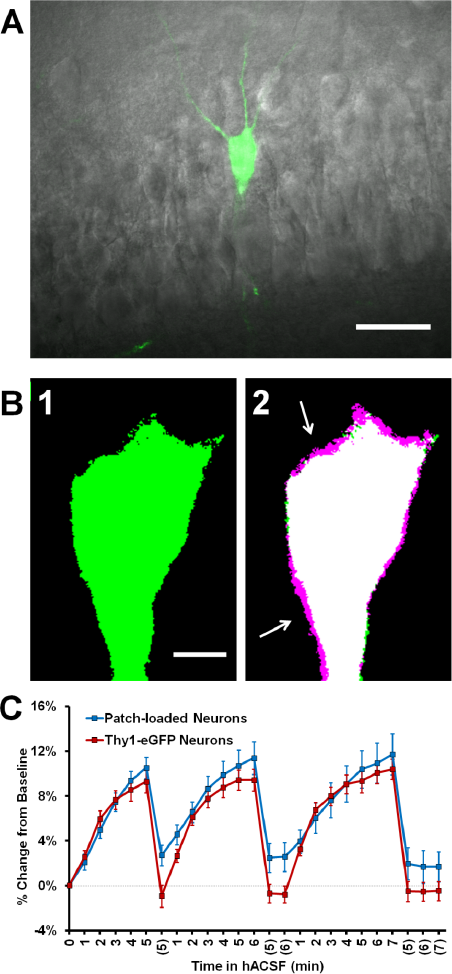
Neuronal swelling persists in Thy1-eGFP neurons. (**A**) Merged infrared (IR) and fluorescent images from a section of the CA1 pyramidal layer in a typical Thy1-eGFP hippocampal slice. GFP-positive neurons are random and generally very sparse, as shown here. (**B**) Representative thresholded images depicting a Thy1-eGFP neuron at standard imaging depth (~30-40 μm deep) at baseline (B1) and 5 minute 40% hACSF time points (B2). Baseline image is overlaid in B2 to reveal soma size increases (magenta regions, indicated by white arrows). (**C**) Volume change in neurons exposed to 40% hACSF, labeled either by patch clamp dialysis of dye ("Patch loaded”, Alexa Fluor 488 or 594 hydrazide) or by expression of eGFP.

**Figure 4:**
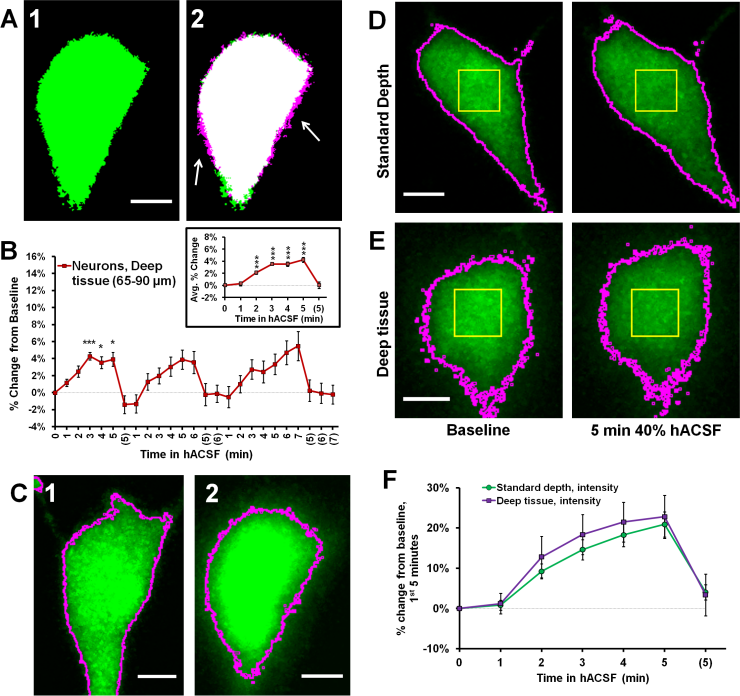
Intensity measurements reveal similar volume change between superficial and deep neurons. (**A**) Representative thresholded images of a Thy1-eGFP neuron taken from deep tissue (~65-90 μm below slice surface) at baseline (A1) and 5 minute 40% hACSF (A2) time points. Magenta regions (indicated by white arrows) show areas of cell volume increase. (**B**) Volume change in deep Thy1-eGFP neurons exposed to 40% hACSF, measured using our standard analysis method. Inset displays average percent change across hACSF applications. The apparent reduction in volume change as compared to shallower neurons (e.g. Figure 3) is most likely an artifact of lower cellular resolution at depth, which is poorly handled by our standard analysis technique. (**C**) Illustration of the analysis problem posed by deeper neurons. Raw image MIPs from 5 minute hACSF time point for a typical shallow (C1) and deep (C2) neuron, illustrating the analysis problem posed by the latter. Images have been brightened to enhance visibility of the cell border. Magenta lines indicate borders of thresholded area for each cell after final processing (see figs. 3B and 4A for respective thresholded images), from which soma area is typically calculated. Shallower neurons (C1) have well-defined borders in the raw MIP which are almost completely preserved after thresholding, allowing for highly-accurate measurements of soma area in these cells. In contrast, the “fuzzy” borders of deeper neurons (C2) are poorly thresholded and insufficient for detecting volume changes. Scale bars = 5 μm. (**D**) Standard (thresholding) and microspectrofluorimetric (fluorescence intensity) methods applied to a standard-depth neuron at baseline (left) and 5 minute 40% hACSF (right) time points. As in (C), magenta outlines indicate cell area calculated by thresholding. Yellow ROIs denote the 5 pm x 5 pm region from which average fluorescence intensity was measured. (**E**) Analysis methods from (D) applied to a deep neuron. While thresholded borders are inaccurate as expected, fluorescence intensity within the yellow ROI decreases to a similar degree between this deep neuron and the more superficial neuron in (D). (**F**) Percent change over the 1^st^ 5 minutes of 40% hACSF, calculated from changes in average fluorescence intensity in both standard depth and deep neurons. ***p < 0.001, *p < 0.05, percent change vs. 0% (baseline).

### 3.3 Neuronal swelling is not due to activation of AMPA receptors or action potential-mediated synaptic transmission

Having ruled out the possibility that neuronal swelling was an artifact of patch clamp, we next considered the specific composition of our solutions as a possible contributor. Until this point, neuronal swelling had been observed in our standard hACSF, which contained 0 mM Mg^2+^, 1 μM TTX and 10 μM NBQX. These conditions are ideal for isolating NMDA receptor-dependent SICs (Lauderdale et al. 2015), but are otherwise quite non-physiological. To determine if the composition of our hACSF was contributing to the observed neuronal swelling, we prepared “antagonist-free” hACSF solutions diluted directly from our standard ACSF (see Methods). Despite the removal of antagonists and partial restoration of Mg^2+^, we continued to observe neuronal swelling across multiple doses of this more “physiological” hACSF (Figure 5A-C). Neuronal swelling was partially dose-dependent (Figure 5D), differing significantly between 17% and 40% hACSF (p = 0.005), but not between 10% and 17% hACSF (p = 0.935). These results provide evidence that neuronal swelling is a direct result of reduced osmolarity rather than an artifact of any particular pharmacological reagent.

**Figure 5:**
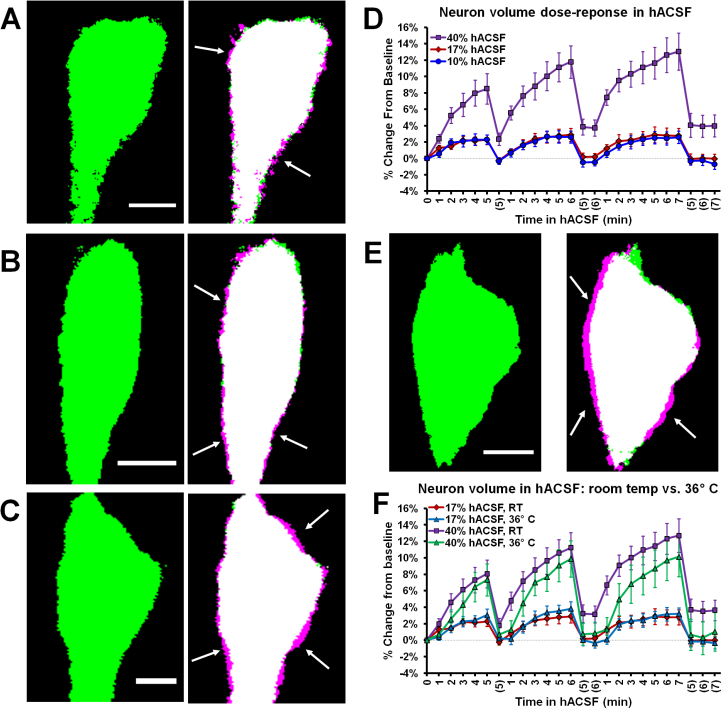
Neuronal swelling occurs across a range of hypoosmolar doses and at physiological temperature. Volume changes observed in Thy1-eGFP neurons exposed to antagonist-free 10% (**A**), 17% (**B**), or 40% hACSF (**C**) at room temperature. Thresholded baseline images (green, left) are overlaid with images taken 5 min. after exposure to hACSF (right) to illustrate swelling of the soma (magenta edges, indicated by white arrows). (**D**) Neuronal volume as percent change from baseline during application of 10%, 17% and 40% antagonist-free hACSF doses. (**E**) Thresholded images of a neuron at baseline (left) and after 5 minutes in 40% hACSF (right), recorded at physiological temperature (36° C). As in A-C, right image is an overlay of 5 minute hACSF (magenta) and baseline (green) time point images, with white arrows indicating magenta regions of cell soma expansion. (**F**) Volume changes compared between neurons imaged at room temperature and 36° C, in both 17% and 40% hACSF. No significant difference was observed between the two temperatures at either dose of hACSF. Scale bars, 5 μm each for A-C and E.

Cellular volume changes and volume regulation can be augmented by higher temperatures (Andrew et al. 1999; Andrew et al. 1997). Therefore, neuronal volume changes might be negligible at physiological temperature due to more efficient volume regulatory mechanisms. To test this possibility, we performed experiments at physiological temperature (~36° C). Neuronal swelling persisted in antagonist-free hACSF at physiological temperature (Figure 5E), and in fact there was no significant difference in neuronal swelling between 22° C and 36° C for either 17% hACSF (F(1,13) = 0.002, p = 0.967) or 40% hACSF doses (F(1,15) = 1.521, p = 0.236; Figure 5F). These results strongly argue against a contribution of temperature to the observed neuronal swelling.

### 3.4 Swelling profiles of layer V cortical pyramidal neurons differ from those of CA1 pyramidal neurons

Recent studies suggest considerable heterogeneity among neurons and astrocytes in different cortical regions, and previous work has indicated that cortical pyramidal neurons (Andrew et al, 2007), but not astrocytes (Risher et al, 2009) may be osmoresistant. To explore the possibility that swelling profiles of astrocytes and neurons are region-specific, we also measured volume changes in pyramidal neurons from neocortical layer V and passive astrocytes from layer II/III (Figure 6). We found eGFP expression to be particularly low in the cortex of Thy1-eGFP mice, which limited their utility for these experiments. Data were therefore pooled from Thy1-eGFP neurons (n = 2) and patch-clamped neurons (n = 8), and compared to pooled hippocampal neurons acquired using both techniques (n = 9 Thy1-eGFP, n = 10 patch clamp). As expected from our observations of hippocampal astrocytes and neurons, cortical layer II/III astrocytes (Figure 6A) and layer V pyramidal neurons (Figure 6B) both exhibited rapid volume changes upon exposure to antagonist-free 40% hACSF. Surprisingly, we observed significantly greater swelling in cortical neurons compared to cortical astrocytes (Figure 6C). Average percent change over 3 hACSF applications (Figure 6D) in cortical neurons was nearly double that of hippocampal neurons (5 minutes hACSF: CA1 neurons, 8 ± 0.81% increase; Cortical neurons, 14 ± 0.98% increase; p < 0.001). The average percent change did not differ between cortical vs. hippocampal astrocytes (F(1,19) = 0.43, p = 0.52; Figure 6E). It was not immediately apparent whether the increase in cortical neuron osmosensitivity reflected an intrinsic difference in membrane properties, or simply a greater freedom for cell swelling due to the looser packing density of neurons in the cortex. These data suggest that osmosensitivity of pyramidal neurons varies by brain region, and importantly, that significant swelling of pyramidal neurons is not restricted to the hippocampal region.

**Figure 6:**
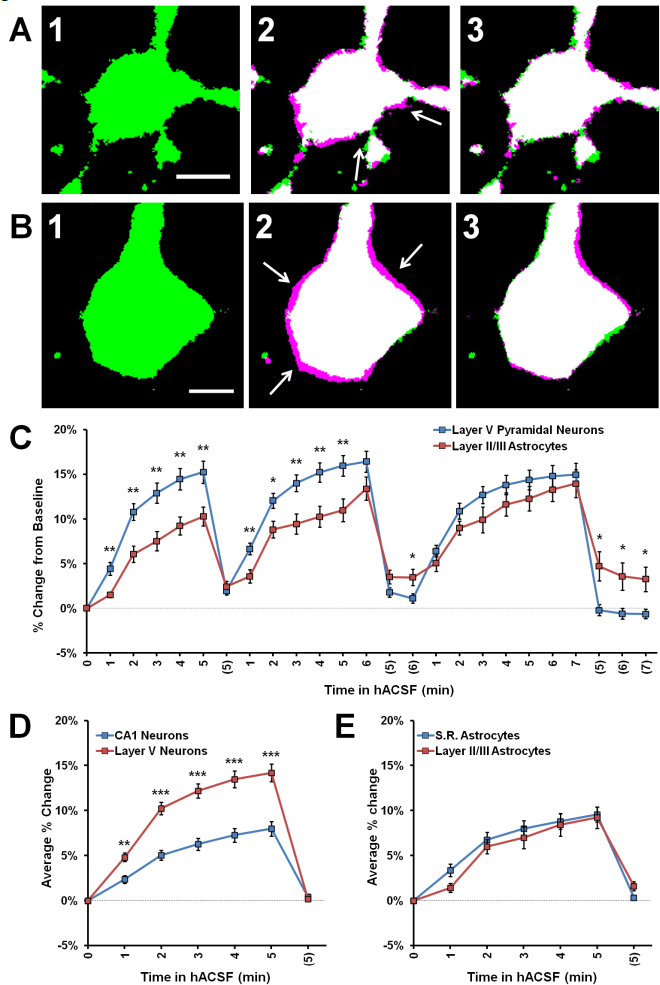
Cortical neurons swell significantly more than cortical astrocytes in 40% hACSF. (**A**) Representative thresholded image of an astrocyte from cortical layer II/III at baseline (A1), overlaid with an image after 5 minutes in antagonist-free 40% hACSF (A2) and after 5 minutes of return to standard ACSF (A3). (**B**) Representative pyramidal neuron from cortical layer V at baseline (B1), overlaid with 5 minute antagonist-free 40% hACSF (B2) and 5 minute standard ACSF wash (B3) time point images, as in (A). Magenta highlights surrounding the cell border in (A2) and (B2), indicated by white arrows, reflect increases in cell volume over baseline. Scale bars, 5 μm. (**C**) Percent change from baseline quantified for cortical (Layer V) neurons and cortical (layer II/III) astrocytes. Note that cortical neurons swell significantly more than astrocytes during the first two applications of hACSF. (**D**) Comparisons of average percent change over the three hACSF applications reveal that cortical neurons swell to approximately twice the size of hippocampal neurons in hACSF, while cortical astrocytes (**E**) show similar responses as hippocampal astrocytes. *p < 0.05, **p < 0.01, ***p < 0.001, layer V neurons versus: layer II/III astrocytes (C) or CA1 neurons (D).

### 3.5 Neuronal swelling is not due to NMDA receptor activation

In examining alternate explanations for neuronal swelling in hACSF, we next chose to more closely investigate the possible involvement of NMDA receptor excitotoxicity. Neuronal swelling is a well-established consequence of excitotoxic NMDA receptor activation (Choi 1992; Liang et al. 2007; Rothman and Olney 1987; Rungta et al. 2015), a distinct possibility in our conditions given the lack of Mg^2+^ in standard hACSF or 40% reduction of Mg^2+^ in physiological hACSF. Additionally, we have previously established that large, NMDA receptor-dependent slow-inward currents (SICs) occur in hACSF (Fiacco et al. 2007; Lauderdale et al. 2015). While SIC frequency (Figure 7A) and amplitude (Figure 7B) can vary between multiple hACSF applications, their rapid appearance during hACSF (Figure 7C) correlates closely with the rapid time course of neuronal swelling and could indicate a common mechanism. To determine whether NMDA receptor activation contributes to neuronal swelling in 40% hACSF, we examined neuronal swelling in our standard hACSF conditions (0 mM Mg^2+^, 1 μM TTX and 10 μM NBQX) with the addition of NMDA receptor inhibitors. Despite the similar circumstances under which SICs and neuronal swelling are observed, we found that neither the selective NMDA receptor antagonist DL-2-Amino-5-phosphonopentanoic acid (DL-AP5; 50 μM), nor 6 mM Mg^2+^, both of which entirely block NMDA receptor-dependent currents (Dissing-Olesen et al. 2014; Lauderdale et al. 2015), had any effect on neuronal swelling in 40% hACSF (F(2,20) = .06, p = 0.942; Figure 7D). These data suggest that neuronal swelling occurs independently of NMDA receptor activation.

**Figure 7:**
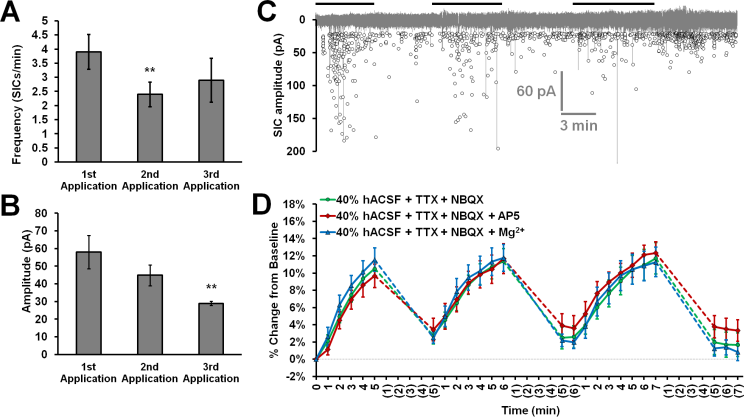
Neuronal swelling is not due to NMDA receptor activation. (**A**) Average SIC frequency and (**B**) amplitude during each application of 40% hACSF; **p < 0.01 versus 1^st^ application. (**C**) Aggregated representation of SIC activity over a typical course of hACSF treatment. Black bars above the trace indicate periods of 40% hACSF. Background trace (grey) shows a voltage-clamp recording of NMDA receptor activity from a typical CA1 pyramidal neuron during 40% hACSF application, while foreground scatterplot shows the overall distribution of SIC activity across all neurons over time (n = 11). (**D**) Neuron soma area measured as percent change from baseline over multiple applications of standard 40% hACSF (containing 10 μM NBQX and 1 μM TTX, to match conditions in which SICs are observed), 40% hACSF + 50 μM AP5, or 40% hACSF + 6 mM Mg^2+^. Time points in (C) and (D) are vertically aligned to facilitate comparison of neuronal SICs and volume changes. Neither AP5 nor high Mg^2+^; were effective in blocking neuronal swelling.

### 3.6 Neuronal and glial swelling occur in tissue from both juvenile and adult mice

We next considered the possibility that neuronal osmosensitivity varied as a function of age. All of our experiments to this point had been performed in slices from P15-P21 juvenile mice. To test the possibility that neuronal swelling was an age-dependent effect, we repeated experiments using adult (10-15 week old) mice (Figure 8). Once again antagonist-free 40% hACSF was prepared, as the inclusion of Mg^2^+ and lack of antagonists provided greater physiological relevance for our results while enabling direct comparison to results from previous studies (Andrew et al. 2007; Risher et al. 2009). As we had observed in juveniles, both adult CA1 neurons and s.r. astrocytes swelled rapidly in response to 40% hACSF application (Figure 8A), and did not differ significantly in their time courses (F(1,17) = 0.61, p = 0.447). Comparison of average percent change between adults and juveniles (Figure 8B, C) revealed no significant differences in neuronal swelling (F(1,16) = 0.09, p = 0.773) or astrocyte swelling (F(1,19) = 0.006, p = 0.94) between the two age groups. These findings suggest that neuronal swelling is not a developmental phenomenon.

**Figure 8:**
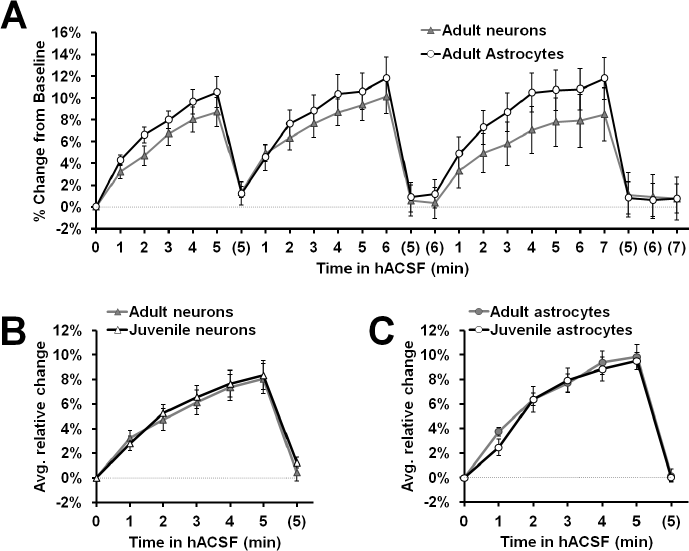
Neuron and astrocyte volume changes in 40% hACSF do not differ between juveniles and adults. (**A**) Rapid neuron and astrocyte swelling was observed in adult hippocampal slices when repeatedly exposed to physiological 40% hACSF. As with juveniles, adult astrocytes and neurons did not differ significantly in their responses to hACSF. (**B**) Average percent change across 3 hACSF applications for adult and juvenile CA1 pyramidal neurons. (**C**) Average percent change across 3 hACSF applications for adult and juvenile s.r. astrocytes. Adults and juveniles did not differ in average percent change for either cell type shown in B and C.

### 3.7 Hypoosmolar swelling of neurons and astrocytes does not require AQP4

In astrocytes, water permeability is thought to be regulated by the expression and gating of aquaporin-4 (AQP4) channels (Gunnarson et al. 2008; Nagelhus et al. 2004; Song and Gunnarson 2012). Functional water channels have not been found in pyramidal neurons, a proposed reason behind their apparent resistance to osmotic stress (Andrew et al. 2007; Gorelick et al. 2006). Our observation of neuronal swelling in all hACSF conditions, however, was clear evidence that the lack of functional aquaporins was not a limiting factor for cell volume change in neurons, and that water must be entering through an alternate route. Several of the alternate, AQP4-independent mechanisms proposed for swelling in astrocytes, including water-permeable cotransporters and even water flux directly across the cell membrane (Kimelberg 2005), could also be functional in neurons and contribute to the observed neuronal swelling in our conditions. If these AQP4-independent mechanisms mediate the rapid neuronal swelling in our hACSF, it follows that the rapid astrocyte swelling in our hACSF may also be partially (or fully) independent of AQP4.

We therefore examined the acute volume responses of both CA1 pyramidal neurons and s.r. astrocytes to antagonist-free hACSF, using hippocampal slices from AQP4^-/-^ mice (Figure 9). As foreshadowed by our neuronal data gathered to this point, astrocytes continued to swell even in the absence of AQP4, implying rapid astrocyte volume changes occur independently of AQP4 expression (Figure 9A). Surprisingly however, AQP4^-/-^ astrocytes swelled significantly more in 40% hACSF compared to wild-type, reaching approximately 1.4 times the volume of wild-type astrocytes by the end of each hACSF application (Figure 9B; AQP4^-/-^, 113.19% ± 0.92%; WT, 109.71% ± 0.54%; p = 0.005). By contrast, astrocyte swelling did not differ between AQP4^-/-^ and wild-type astrocytes in 17% hACSF (F(1,17) = 1.000, p = 0.331). We also tested the effects of AQP4^-/-^ on neuronal volume changes in 40% hACSF. Not surprisingly, AQP4 deletion had no effect on neuronal swelling (F(1,16) = 0.02, p = 0.891; Figure 9C, D). These data seemed to indicate that AQP4 expression, far from being *required* for intracellular accumulation of water in astrocytes, may actually play an important role in *limiting* the degree of swelling during severe hypoosmolar stress. Overall, our results strongly suggest that water influx resulting from hypoosmolar ACSF application is not selective to astrocytes, and is not mediated by AQP4.

**Figure 9:**
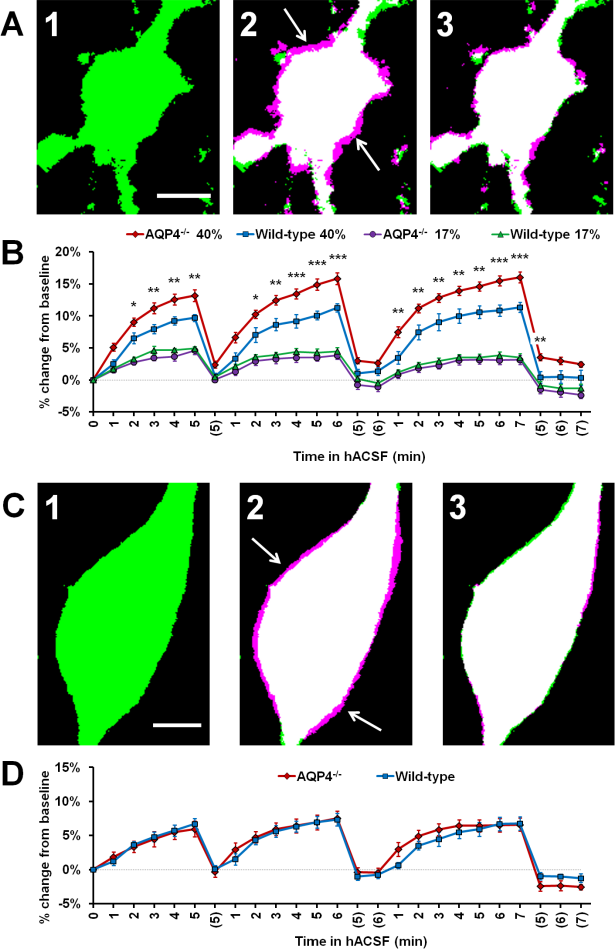
Astrocyte volume changes do not require expression of AQP4. (**A**) Representative thresholded images of an AQP4^-/-^ astrocyte soma at baseline (A1), overlaid with an image after 5 minutes in antagonist-free 40% hACSF (A2) and after 5 minute wash in standard ACSF (A3). As in previous figures, magenta regions (indicated by white arrows) represent increases in volume over baseline. (**B**) Astrocyte volume, quantified as percent change from baseline, in AQP4^-/-^ and wild-type astrocytes exposed to 17% or 40% hACSF. AQP4^-/-^ astrocytes showed no change in swelling at 17% hACSF and significantly greater swelling at 40% hACSF, as compared to wild-type. (**C**) Representative thresholded images and overlays of an AQP4^-/-^ neuron soma at baseline (C1), after 5 minutes in antagonist-free 40% hACSF (C2), and after a 5 minute wash period in standard ACSF (C3). (**D**) Volume changes in hACSF are compared between AQP4^-/-^ and wild-type neurons. As expected, neuronal swelling did not differ significantly between these two genotypes. *p < 0.05, **p < 0.01, and **p < 0.001 versus wild-type astrocytes in 40% hACSF. Scale bars, 5 μm each for (A) and (C).

## 4 Discussion

The major finding in this study is that CA1 pyramidal neurons readily swell in hypoosmolar conditions, increasing their volume significantly above baseline even within the first minute of hACSF application. The rate and overall amount of neuronal swelling was found to be almost identical to that observed in stratum radiatum astrocytes, and was observed repeatedly across a variety of experimental conditions. Neuronal swelling was rapidly reversible, returning to baseline after 5-7 minutes in normosmolar ACSF, and readily returned upon re-application of hACSF. While the astrocyte swelling observed is in line with the reportedly high osmosensitivity of astrocytes and astrocyte-like glial cells (Hirrlinger et al. 2008; Risher et al. 2009), the neuronal swelling observed contrasts with previous work concluding that neurons steadfastly retain their volume during osmotic challenge (Andrew et al. 2007; Caspi et al. 2009). Neither glial nor neuronal swelling was found to require AQP4 expression.

These data cast doubt over the commonly-accepted roles of neurons and astrocytes in brain water regulation.

The possibility that neuronal volume changes are extremely sensitive to subtle differences in methodology was strongly considered and systematically examined in this study. Initial experiments were performed in tissue slices from juvenile mice at room temperature, using whole-cell patch clamp to dialyze individual CA1 hippocampal cells located 30-50 μm from the slice surface with fluorescent dye. These conditions differed substantially from those of Andrew et al. (2007), the study most widely cited as concrete evidence of neuronal osmoresistance. For example, Andrew and coworkers performed experiments at 33-34°C based on a previous observation that slice swelling is noticeably decreased at higher temperatures (Andrew et al. 1997), a possible indicator of isovolumetric regulation (IVR) which is characterized by an apparent resistance to volume change (Franco et al. 2000; Lohr and Grantham 1986; Pasantes-Morales and Tuz 2006; Tuz et al. 2001). Furthermore, isovolumetric regulation is observed only during gradual (but not sudden) shifts in osmolarity (Franco et al. 2000; Pasantes-Morales and Tuz 2006; Tuz et al. 2001). Neurons imaged deeper in tissue would likely be exposed to more gradual solution exchanges and therefore, more gradual changes in osmolarity which could reduce the rate of swelling and facilitate IVR. Therefore, to specifically test the role of these and other variables, additional recordings were performed in the following conditions: 1) Using Thy 1-eGFP transgenic mice (Andrew et al. 2007; Feng et al. 2000) to avoid possible cell damage from patch clamp; 2) At physiological temperature (36°C); 3) In deep tissue (65-90 μm); 4) In slices from adult (10-15 week-old) mice; and, 5) In the neocortex. Neuronal swelling was observed across all of these conditions, suggesting that it is a ubiquitous response to reductions in osmolarity in both juvenile and adult tissue.

Using our standard volume analysis protocol, we found that swelling of neurons located deeper in the tissue (65-90 jm) was more difficult to observe compared to superficial neurons. We interpret much of this apparent difference as a loss of image fidelity rather than a true effect of tissue depth on volume changes. This interpretation is supported by our observation that changes in cell fluorescence intensity over time (which are inversely related to changes in cell volume; Crowe et al. 1995) did not significantly differ between superficial and deep neurons-a strong indication that neurons at both depths were swelling to similar volumes.

It is worth noting that our standard image analysis protocol, although based on established methods (Hirrlinger et al. 2008; Risher et al. 2009; Risher et al. 2012), was intentionally conservative and likely underestimates actual changes in cell volume. Cell soma area was chosen as a proxy for overall volume changes, since direct measures of cell volume were not feasible for our conditions. Assuming that cell volume is changing equally in all directions, it can be inferred that our measurements reflect only about two-thirds of the true change in volume. This inference is partially supported by the analysis of fluorescence intensity as an indication of cell volume changes, which suggested volume increases > 1.5 times those calculated from thresholded images. We also observed that the precise method of image processing had a surprisingly strong impact on measured volume. The overall magnitude of volume change observed could often be increased twofold or more simply through the use of different thresholding methods. These apparent increases in signal-to-noise ratio often came at the expense of “cell” and “background” pixels being incorrectly classified, and the thresholded images inconsistent with what could be observed “by eye” pre-thresholding (unpublished observations). We therefore rejected these methods in favor of the more conservative “mean” thresholding method, which more consistently returned an image in which the cell body could be clearly identified.

One of the more exciting findings in the present study was a lack of evidence for AQP4-mediated water *influx* into astrocytes or neurons exposed to hACSF. While not necessarily surprising for neurons, which do not express AQP4 (Nielsen et al. 1997), lack of inhibition of astrocyte swelling in AQP4^-/-^ mouse tissue was rather unexpected based on a large body of literature on the requirement of AQP4 for facilitated water movement across the astrocyte membrane. Surprisingly, not only was AQP4 not required for astrocyte swelling, but astrocyte volume in severely (40%) hypoosmolar conditions was significantly greater in AQP4^-/-^ mice compared to wild-type animals. There are a couple of potential explanations for this finding.

First, AQP4^-/-^ animals have been shown to have a nearly 30% larger resting brain extracellular space compared to wild-type animals (Binder et al. 2004; Yao et al. 2008). It is possible that the enlarged ECS in AQP4^-/-^ mice provides more available space for astrocytes to swell into, leading to increased astrocyte swelling. This explanation, however, fails to account for the lack of increased neuronal swelling in AQP4^-/-^ mice, since neurons would presumably also be adjacent to increased extracellular space. It is also unlikely that astrocyte volume changes in our 40% hACSF were limited by the extracellular space in wild-type animals, as tissue continues to swell in far more severe hypoosmolar conditions (Chebabo et al. 1995b).

A second explanation is that AQP4 can provide an efficient efflux pathway for water to leave astrocytes. AQP4 expression is polarized, with highest levels of expression in perivascular endfeet (Nielsen et al. 1997). Loss of AQP4 at endfeet has been proposed to inhibit water removal from the astrocyte, leading to increased swelling (Wetherington et al. 2008) and impairing tissue recovery from vasogenic edema (Papadopoulos et al. 2004). Recent work demonstrates AQP4-dependent water efflux may also be important physiologically. Haj-Yasein and colleagues observed that Schaffer collateral stimulation in AQP4^-/-^ hippocampal slices resulted in a greater ECS reduction (indicating increased cell swelling), and higher peak [K+]_o_ compared to slices from wild-type mice (Haj-Yasein et al. 2015; Haj-Yasein et al. 2012). The authors concluded that ECS shrinkage (caused by activity-dependent astrocyte water uptake) is normally offset by an osmotically-driven water efflux through perisynaptic AQP4 channels. Our findings suggest that bath application of hACSF into the slice similarly produces a rapid AQP4-independent influx of water into the astrocyte, from which it cannot efficiently escape because the AQP4 efflux pathway has been removed.

Interestingly, some recent studies suggest that rapid water influx through AQP4 may be important for opening stretch activated channels associated with astrocyte volume regulation (Benfenati et al. 2011; Jo et al. 2015; Mola et al. 2016). This so-called “isovolumetric regulation” (IVR), involving efflux of intracellular osmolytes into the ECS, is generally observed with small and gradual changes in osmolarity, and in some cases can completely prevent swelling of glial cells or tissue slices (Franco et al. 2000; Lohr and Yohe 2000). IVR is easily overwhelmed by more rapid or larger osmotic shifts (Lohr and Yohe 2000), which may explain the lack of visible IVR in the current study. Thus, it is possible that volume regulatory mechanisms are actively resisting volume change in wild-type astrocytes, and that such mechanisms are impaired in AQP4^-/-^ astrocytes. Regardless of underlying mechanism, our findings suggest that: a) AQP4 is not required for rapid water influx into CA1 neurons or astrocytes; and b) AQP4 may serve as an important water *efflux* pathway that helps limit the extent of astrocyte swelling.

The observation that the effects of AQP4^-/-^ on astrocyte swelling are hACSF dose-dependent is not unprecedented. Thrane et al. (2011) observed a reduction in AQP4^-/-^ astrocyte swelling in 20% hACSF, but no difference in 30% hACSF. A direct comparison of their findings to ours is somewhat complicated, as Thrane et al. used different hACSF doses and longer imaging intervals (5-10 min) relative to ours. Based on the direction of the effect of AQP4 deletion on astrocyte swelling, however, it is tempting to speculate that Thrane and coworkers might have also observed augmented astrocyte swelling in 40% hACSF. It is also important to note that Thrane et al. measured AQP4^-/-^astrocyte volume in the cortex, while we recorded effects on s.r. astrocytes in CA1. AQP4 expression levels and patterns are quite distinct between these two regions (Hubbard et al. 2015), and it is possible that the role of AQP4 in water movement is also brain region-dependent. Although the exact effects of AQP4 deletion on astrocyte swelling differed between the two sets of data, both suggest that the role of AQP4 in water movement is governed by the severity of the osmotic challenge.

If neurons are indeed swelling in hACSF, then what is the route of water influx? Clues to this answer may lie in other conditions which cause neuronal swelling. While hypoosmolar swelling has generally been reported only in dissociated neurons *in vitro* (Aitken et al. 1998; Boss et al. 2013; Inoue et al. 2005; Somjen 1999), pyramidal neurons have long been known to swell in excitotoxic conditions (Choi 1992; Risher et al. 2009; Zhou et al. 2010). Massive intracellular sodium influx (as observed in cases of continuous neuronal firing or excessive NMDA receptor activation) can cause sufficient depolarization to open voltage-gated Cl^-^ channels, with the resulting increase in intracellular osmolarity (from both Na^+^ and Cl^-^) producing an osmotic gradient into the neuron (Rungta et al. 2015). This highlights the fact that, despite lacking functional water channels, neurons are capable of taking on water through alternate means quite rapidly. Consistent with this, spreading depression (a wave of depolarization, excessively high K+ and glutamate release in the extracellular space) has been reported in hypoosmolar conditions (Chebabo et al. 1995a; Huang et al. 1995) and can induce excitotoxic neuronal swelling (Zhou et al. 2010). There is little reason to believe an excitotoxic mechanism is directly responsible for the observed neuronal swelling in our model, as it was unaffected by NMDA receptor blockers, and we have previously found no evidence of spreading depression in hACSF (Lauderdale et al. 2015). Instead, we propose that the osmotic gradient, which drives water into a neuron following excessive Na^+^ and Cl^-^ influx (Rungta et al. 2015), parallels the gradient imposed by hACSF on a neuron with normal intracellular ion concentrations. Routes for such water influx into neurons are not well defined. Two possible routes might be inferred from the known or hypothesized AQP4-independent pathways for water influx into astrocytes: simple water diffusion across the plasma membrane (Kimelberg 2005; Papadopoulos and Verkman 2013), or water movement through a commonly-expressed cotransporter such as the Na^+^,K^+^,2Cl^-^ cotransporter NKCC1 (Jayakumar and Norenberg 2010; Kimelberg 2005; Macaulay and Zeuthen 2012; MacVicar et al. 2002; Su et al. 2002a; Su et al. 2002b; Yan et al. 2003). Neither of these water pathways is likely to be exclusive to astrocytes, and the similarity in swelling profiles between neurons and astrocytes in our standard hACSF conditions might suggest a common mechanism independent of AQP4. Future studies will explore potential mechanisms of water influx into astrocytes and neurons in hypoosmolar conditions.

